# GWAS reveal a role for the central nervous system in regulating weight and weight change in response to exercise

**DOI:** 10.1101/2020.09.10.291229

**Authors:** Louis P. Watanabe, Nicole C. Riddle

## Abstract

Body size and weight show considerable variation both within and between species. This variation is controlled in part by genetics, but also strongly influenced by environmental factors including diet and the level of activity experienced by the individual. Due to the increasing obesity epidemic in much of the world, there is considerable interest in the genetic factors that control body weight and how weight changes in response to exercise treatments. Here, we use GWAS in Drosophila to identify the molecular pathways that control weight and exercise-induced weight changes. We find that there is a complex set of molecular pathways controlling weight, with many genes linked to the central nervous system (CNS). Weight was strongly impacted by animal size and body composition. While the CNS appears to be important for weight and exercise-induced weight change, signaling pathways are particularly important for determining how exercise impacts weight.

## INTRODUCTION

Comparing individuals of different species, it is clear that body size is a species-specific characteristic. However, within species a range of body sizes and weights typically can be observed. At least some of this variation is due to genetic factors which impact body size (Boulan et al., 2015; Flatt, 2020; Jimenez, 2016; Mirth & Shingleton, 2012). This impact of genetics is illustrated by dwarf mutations that have been observed in a variety of species, including plants and animals [for example, see (Ferrero-Serrano et al., 2019; Jaarsma et al., 2013)]. For example, mutations in hormone production or signaling pathways often cause body sizes outside of the normal range for the species, be that dwarfism or gigantism (Bartke & Quainoo, 2018; Beckers et al., 2018). Environmental factors also play an important role in determining body size and weight, as illustrated by the typically smaller size of individuals in harsh environments such as mountain tops or the larger size reached by individuals in nutrient-rich milieus (Hyun, 2013; Koyama et al., 2020; Oldroyd & Leyser, 2020; Werner & Schmulling, 2009). In addition, the rising obesity rates in countries across the world during the last few decades well illustrate the impact of environmental factors on body size and weight, as the largest contributors to this obesity epidemic are identified as inexpensive, high-energy diets coupled with increasingly sedentary lifestyles (Hruby & Hu, 2015; Lifshitz & Lifshitz, 2014; Seidell & Halberstadt, 2015). Thus, a combination of genetic and environmental factors as well as their interaction determine an individual’s body size and weight.

Given the increasing incidence of obesity worldwide, there is growing interest in understanding the genetic architecture controlling body size and weight. Also of strong interest is the role of genetics in an individual’s response to exercise, which is one of two primary treatments for obesity (the other being modification of diet). Because exercise has many benefits and typically involves low risks, many medical professionals recommend exercise as part of an individual’s weight-loss regime (Donaldson et al., 2015; Kahan & Zvenyach, 2016). Although there is considerable variation between individuals, the effectiveness of exercise as part of a treatment program for obesity in general is well documented (Laskowski, 2012; Thomas et al., 2013; Villareal et al., 2017). Despite the importance of exercise as a tool to combat obesity, there are currently no methods to predict how an individual will respond to an exercise treatment because the impact of exercise in different genetic backgrounds is poorly understood (Bouchard, 2011). Thus, research into the genetic factors controlling weight and exercise-induced weight change is needed urgently to improve and personalize exercise treatments for obesity and other conditions.

Model organisms such as *Drosophila melanogaster* and *Caenorhabditis elegans* provide a unique opportunity to further our understanding of the genetic factors controlling body weight, body size, as well as the impact of exercise on weight and other aspects of physiology (Laranjeiro et al., 2017; Riddle, 2019; Sujkowski & Wessells, 2018; Watanabe & Riddle, 2019). While fruit flies and worms have not been used traditionally in exercise research, exercise methods that can complement human and mouse studies have been established within the last two decades for both flies (Mendez et al., 2016; Piazza et al., 2009; Sujkowski et al., 2015; Sujkowski et al., 2017; Tinkerhess et al., 2012; Watanabe & Riddle, 2017, 2018) and worms (Chuang et al., 2016; Hartman et al., 2018; Laranjeiro et al., 2017; Laranjeiro et al., 2019). Exercise systems for Drosophila include the Power Tower, which induces high intensity exercise by repeatedly dropping the fly enclosures, causing flies to drop to the bottom of the vials, and then climbing up due to their innate negative geotaxis (Piazza et al., 2009). The TreadWheel Drosophila exercise system and its cousin the REQS (Rotating Exercise Quantification System) use rotation of the fly enclosures to induce a more gentle, low impact form of exercise (Mendez et al., 2016; Watanabe & Riddle, 2017, 2018). These examples demonstrate that tools to implement exercise treatments in invertebrate model systems are available for scientists to apply them to questions related to exercise physiology and response.

The potential of using these innovative exercise systems in a genetic model such as Drosophila is illustrated by the genome-wide association studies (GWASs) carried out by our laboratory. We utilized the REQS and the Drosophila Genetics Reference Panel (DGRP), a set of approximately 200 fully sequenced, wildtype inbred lines with an established computational GWAS pipeline (Huang et al., 2014; Mackay et al., 2012). Animal activity levels with and without rotational exercise stimulation were measured to gain insights into the genetic architecture controlling these traits (Watanabe et al., 2020). In the GWASs, we identified over 100 genes linked to basal or exercise-induced activity levels in Drosophila, many of which were linked to functions in the central nervous system (CNS) as well as a set of chromatin modifiers with no previous link to animal activity (Watanabe et al., 2020). With many of the genes identified having no reported link to animal activity, this dataset represents a resource for biologists interested in animal activity and exercise and provides outstanding opportunities for working with undergraduate students, as the assays involved are simple and mutants are available readily from stock centers. Thus, this study illustrates how research in model systems can further our understanding of exercise and physiological responses to exercise such as weight change.

Here, we follow up on this activity study, using a similar approach to investigate the genetic factors impacting animal weight and exercise-induced weight change in *Drosophila melanogaster*. We used the DGRP strain collection to carry out a series of GWASs to identify genetic variation linked to animal weight and exercise-induced weight change. We found that there is a large amount of variation in animal weight in this strain collection, variation which is strongly influenced by sex and genotype and under genetic control. Weight change induced by a five-day exercise treatment also was highly variable, including both weight gain and weight loss, with some animals mostly lacking a response. The GWASs revealed a complex genetic architecture underlying the variation in animal weight, identifying approximately 100 candidate genes and suggesting an important role for the CNS in regulating weight. Follow-up studies suggest that the overall variation in animal weight is due to two separate factors, animal size and body composition. Our results also indicate that signaling pathways are particularly important for exercise-induced weight change, which was linked to 52 genes. In females, there was a clear relationship between the amount of exercise performed and the weight change observed. Together, our results demonstrate that both weight and exercise-induced weight change are impacted by a variety of molecular pathways and hint at the importance of long-range communication of the CNS with the body.

## RESULTS

### Animal weight in the DGRP collection is highly variable

To understand how exercise impacts weight, we utilized the Drosophila model system and compared weight of animals having completed a five-day exercise intervention to weight of untreated control animals (see Figure 1 for an experiment overview). 38 genetically distinct strains from the DGRP collection of wild-derived Drosophila strains (Huang et al., 2014; Mackay et al., 2012) were included in this study. The exercise treatment utilized the TreadWheel (Mendez et al., 2016), which uses rotation of fly vials to stimulate increased activity in the animals (Watanabe & Riddle, 2017). Flies in the treatment group were exercised for two hours per day for five consecutive days; untreated control animals were handled the same as the treated flies, but placed next to the TreadWheel in the incubator. The day following the last exercise treatment, both exercise-treated and control animals underwent a fitness test and were weighed in groups of ten flies. We found a large amount of variation in animal weight, both in exercised and unexercised animals (Figure 2). Excluding measurements based on less than three replicates, the lowest weights for all groups, male, female, treated, or untreated, were found in line 324, with control males weighing 0.502 +/− 0.017mg, exercised males weighing 0.516 +/− 0.017mg, control females weighing 0.750 +/− 0.019mg, and exercised females weighing 0.705 +/− 0.018mg. The heaviest flies came mostly from line 310, with control males weighing 0.860 +/− 0.013mg, control females weighing 1.306 +/− 0.011mg, and exercised females weighing 1.337 +/− 0.025mg. The heaviest exercise treated male flies came from line 730, with 0.854 +/− 0.012mg. Thus, the heaviest flies were approximately 2.7 times the weight of the lightest, with the heaviest males weighing ~ 1.7 times the weight of the lightest males, and the heaviest females weighing ~ 1.9 times the weight of the lightest females (Figure 2). We conclude that fly weight varies considerably in the DGRP strain collection and that it might be possible to identify genetic factors controlling weight by GWAS.

**Figure 1.**
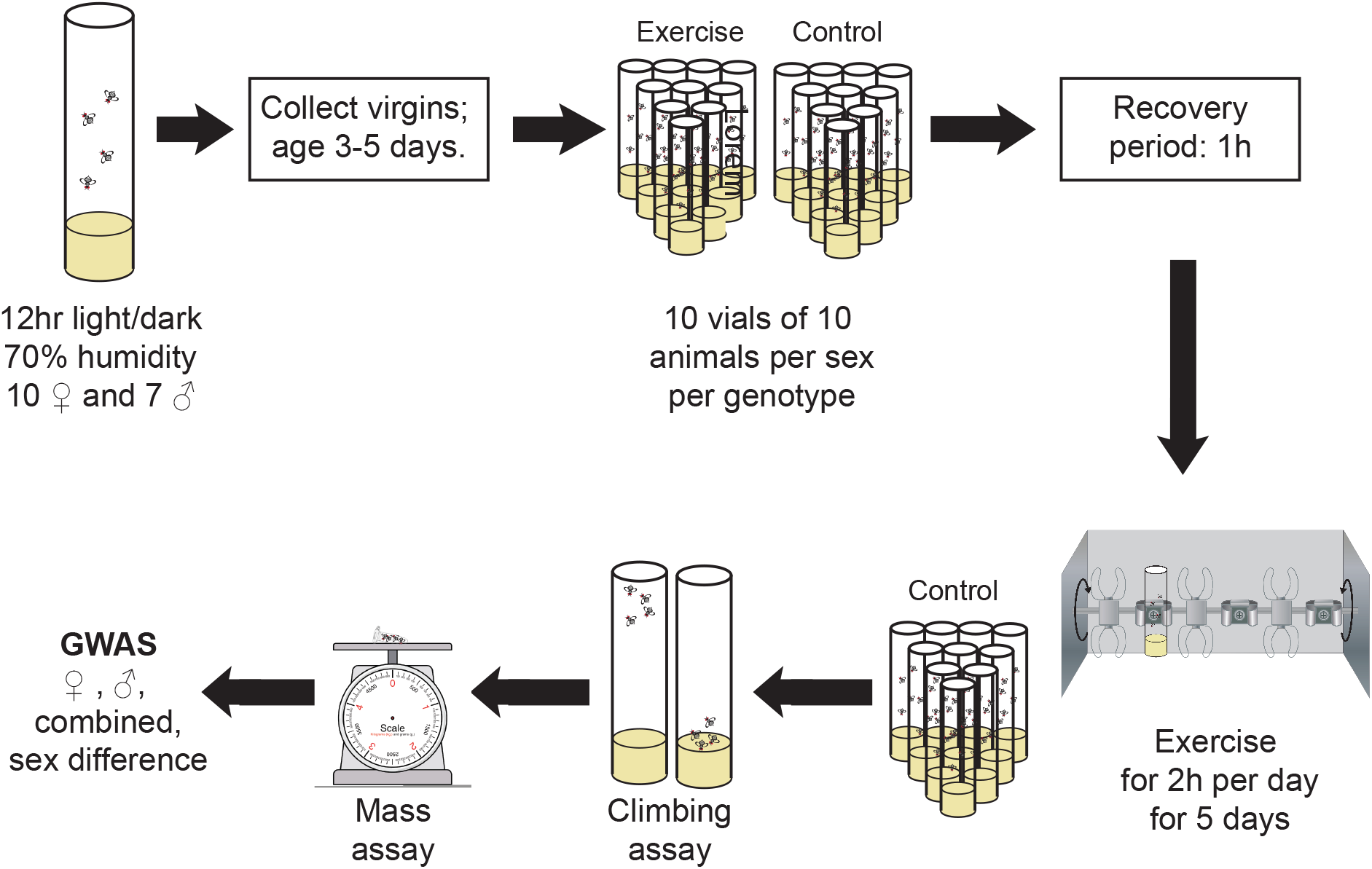
Overview of the experiment. This diagram provides an overview of the study discussed here, illustrating the handling of both control (non-exercised) and treated (exercised) animals.

**Figure 2.**
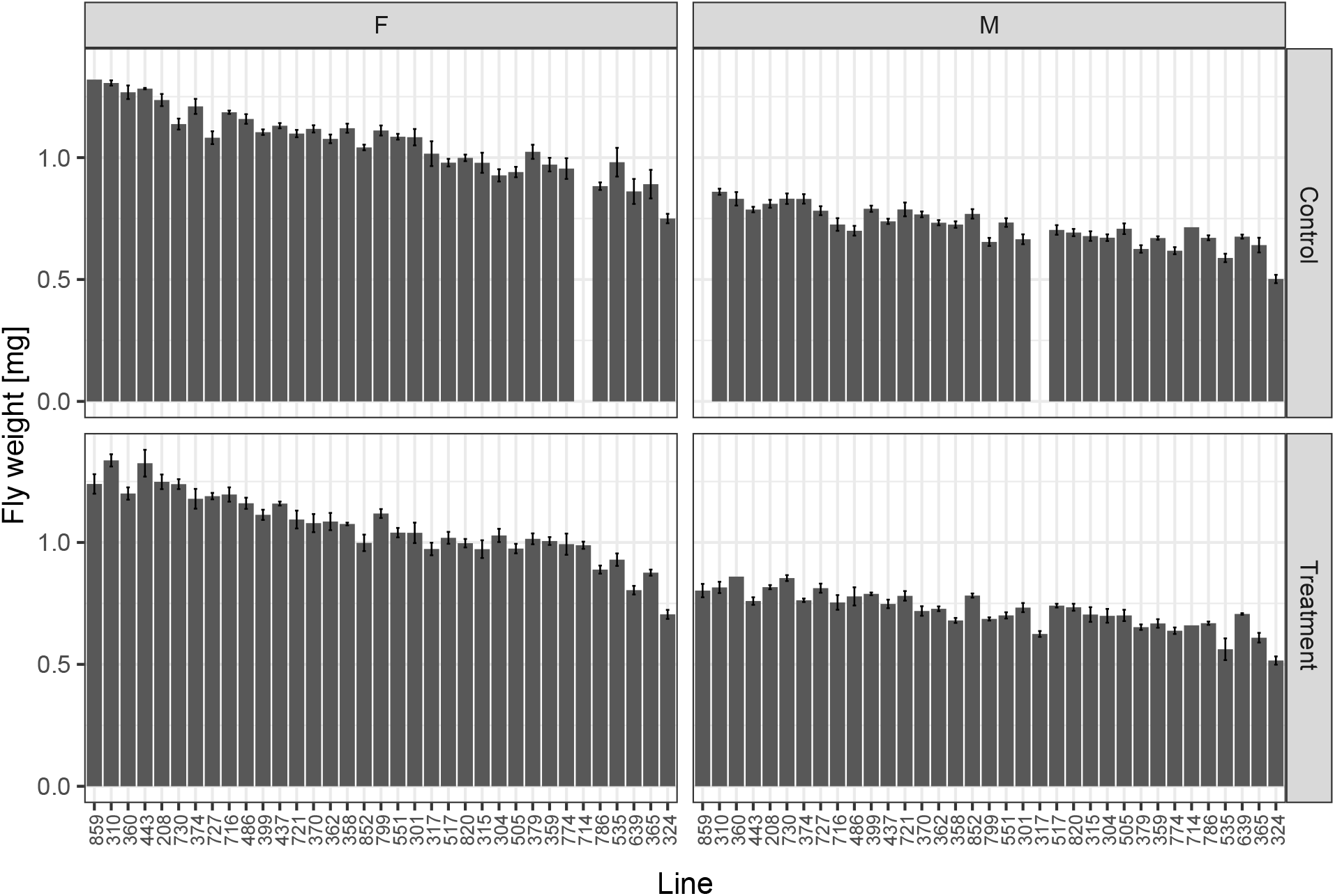
Weights of flies in the DGRP population vary considerably, both with and without exercise treatment. Bar graphs showing group means +/− standard error for weights per fly in mg (Y-axis). The lines are arranged from the highest to lowest overall weight. Top left - data from female (F) control animals; top right – data from male (M) control animals; bottom left – data from female exercise-treated animals; bottom right – data from male exercise-treated animals. Missing data for a specific line/sex/ treatment combination results in missing bars in the graph.

Before investigating possible genetic factors controlling animal weight, we first explored how sex, genotype, and treatment relate to animal weight using Kruskal-Wallis rank sum tests. We found that both sex and genotype (DGRP line) strongly impact animal weight, with p-values highly significant for both sex and genotype (p < 2.2e-16). We found no overall, consistent impact of the exercise treatment on animal weights in this analysis (p = 0.8395), suggesting that any impacts of treatment likely differ between strains and possibly between sexes. Thus, both sex and genotype are significant factors in determining an animal’s weight, supporting the observation that females typically are larger than males, and that animal size is reported to be impacted by a variety of mutations (FlyBase, FB2020_03) (FlyBase et al., 2004; Gramates et al., 2017; Thurmond et al., 2019).

### Animal weight is highly correlated between animals of one genotype, but shows no relationship to activity levels or lifespan

Next, we investigated the relationship between weights in the two sexes and among the two treatment groups (Figure 3). Looking first at the correlation between the two treatment groups, we found that the weights of control and exercise-treated animals were highly correlated, with a correlation of 0.94 in females (Pearson’s product-moment correlation, p < 2.2e-16) and 0.90 in males (p = 1.091e-12). Comparing the weights between sexes, the correlation was 0.80, both in the control animals (p = 2.947e-08) and in exercise-treated animals (p = 9.42e-09). These values suggest that while males and females differ significantly in their weight and size, male and female weights of a specific genotype are strongly correlated with each other, both in the presence and absence of an exercise treatment.

**Figure 3.**
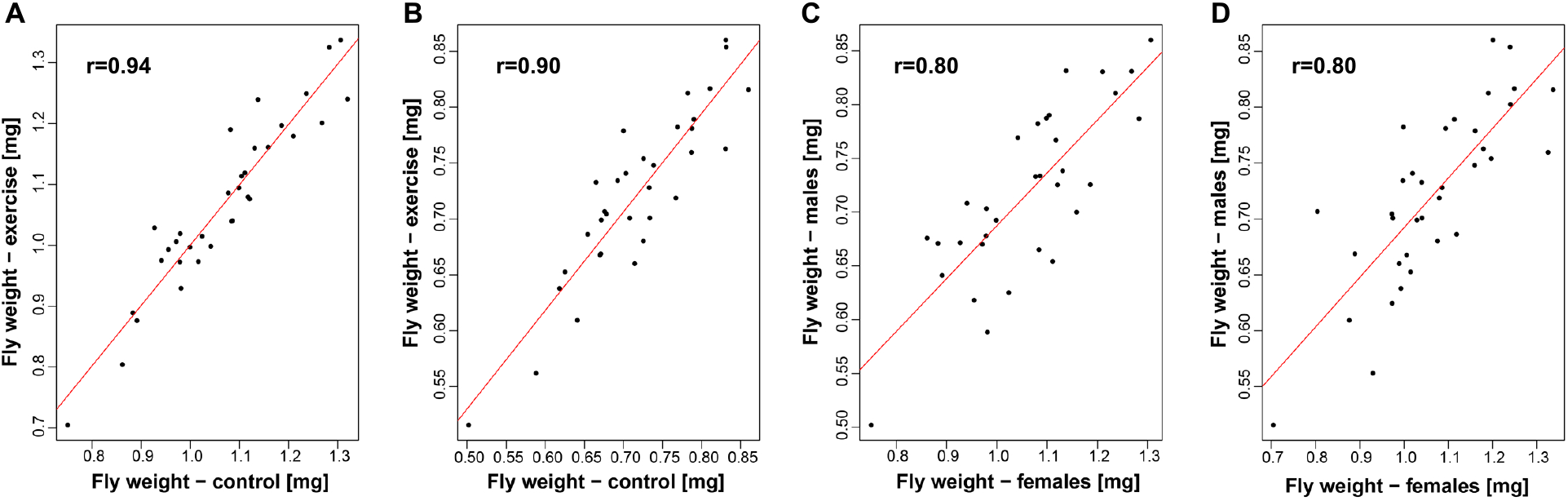
Animal weights are strongly correlated between male and females of the same strain, irrespective of exercise. X- and Y-axis: Weight per fly in mg. r: Pearson’s product-moment correlation. **A.** Data from females; X-axis – control animals; Y-axis – exercise-treated animals. **B.** Data from males; X-axis – control animals; Y-axis – exercise-treated animals. **C.** Data from control animals; X-axis – females; Y-axis – males. **D.** Data from exercise-treated animals; X-axis – females; Y-axis – males.

To gain further insights into the consequences of the weight variation observed in the DGRP strain collection, we investigated if the base weight of the animals was related to the typical activity level of the animals and to their lifespan. We included lifespan in these analyses due to the observation in other species that smaller dwarf animals often show increased lifespan (Bartke, 2017). We found that there is no relationship between baseline activity levels as reported previously by our lab (Watanabe et al., 2020) and weight in this strain collection, neither in females (r = 0.1958404; p-value = 0.3276), nor in males (r = 0.252795; p-value = 0.2128; Figure S1). Based on publicly available lifespan data from females (Durham et al., 2014; Ivanov et al., 2015), there was also no clear relationship between animal weights and lifespan in this study population (data from Durham and colleagues: r = −0.2184477; p-value = 0.2378; data from Ivanov et al: r= −0.1667798; p-value = 0.3616; Figure S2). Thus, while data from other study systems suggests connections between basal animal activity levels, weight, and lifespan, such connections are not detected in this Drosophila population.

### GWASs identify over 100 genetic variants linked to weight in control and exercise-treated animals

As the exploratory analysis revealed the importance of animal genotype for fly weight, we carried out a GWAS for the weight phenotypes measured in our study (weight in the control animals and weight in the exercise-treated animals). To determine if a GWAS was a valid approach for these phenotypes, we calculated heritability as well as a variety of additional quantitative genetics parameters (Table S2). We found that heritability (H^2^) for weight in the control animals was 0.773 and 0.758 in exercise-treated animals. This finding is considerably higher than the heritability of 0.25 reported by Jumbo-Lucioni and colleagues using a similar set of fly lines (Jumbo-Lucioni et al., 2010), but is consistent with studies from other animals that suggest a high heritability for animal weight (and body mass index in humans) (Dubois et al., 2012). This level of heritability confirms that the animal weight data presented here is appropriate for analysis by GWAS.

To elucidate the gene networks controlling animal weight in both control and exercise-treated flies, two separate GWASs were performed, one using the data from control animals, one using the data from treated animals. The analysis was carried out using the webtool developed by Mackay and colleagues for use with the DGRP population (Huang et al., 2014; Mackay et al., 2012). This GWAS tool implements four different analyses if data from both sexes are available, analyses using the combined data, male data, female data, and an analysis investigating the difference between the sexes. The GWAS for weight in the control animals identified 44 genetic variants representing 23 genes in the four analyses (Table S3). Two of the genes identified, *zip* and *lncRNA:CR32773*, had low p-values in the analysis of the combined data as well as the male and female datasets, indicating that these genes might function similarly in both sexes. Among the 23 genes linked to animal weight in this analysis were five genes that are described as “size defective” on FlyBase (FB2020_03) (FlyBase et al., 2004; Gramates et al., 2017; Thurmond et al., 2019): *lncRNA:CR32773, Grip75, Lar, Ptp61F,* and *sfl.* Based on the information available, the most promising candidates among these are *lncRNA:CR32773*, as knockdown of this lncRNA leads to small wing size and is linked to the TGFβ signaling pathway (Hevia et al., 2017); *sfl*, as overexpression of this gene in the wing leads to smaller wing size (Kamimura et al., 2011); and the tyrosine phosphatase *Ptp61F*, where again, overexpression in the wing leads to smaller wing size (Buszard et al., 2013). Thus, the available information about the genes linked by the GWAS to fly weight support their investigation as candidates in further studies.

The GWASs using the weight of exercised animals were carried out as described in the previous paragraph for the weight of the control animals. These analyses identified 179 genetic variants linked to weight after exercise treatment, representing 75 genes (Table S3). There were 18 genes that had low p-values in the male, female, and combined analyses, *CG7560, CG33995, CG43060, CG44153, Lrt, lilli, Lar, lncRNA:CR32773, lncRNA:iab8, Mcm7, mthl15, Nrk, sfl, ssp3, TER94, uzip, Wnt4,* and *zip*, suggesting that these genes impact weight in exercised animals of both sexes. Similar to what was observed for the genes linked to weight in the control animals, we found 13 genes that are described as “size defective” on FlyBase (FB2020_03) (FlyBase et al., 2004; Gramates et al., 2017; Thurmond et al., 2019): *ASPP, Lar, lncRNA:CR32773, lilli, MCU, orai, Ptp61F, pum, Ref1, sfl, tai,* and *uif*. Among this gene set were four “size defective” candidate genes also detected in the analysis of body weight in the control animals, including the three most promising candidate genes, *lncRNA:CR32773, Ptp61F, and sfl* (fourth gene: *Lar*). Among the nine remaining genes, there were several other candidates worth noting. *uif* mutant larvae show significantly reduced body size (they do not survive to the adult stage) (Zhang & Ward, 2009). *orai* mutant animals are smaller than control animals raised under the same conditions (Pathak et al., 2015; Pathak et al., 2017), as are mutants of *tai*, which also show reduced wing size (Zhang et al., 2015). In contrast to the size reduction seen in these mutants, *ASPP* mutants are reported to have increased wing size, suggesting that their body size might be increased as well (Bertran et al., 2019). Together, these data suggest that the gene variants linked to weight in exercise-treated animals include several linked to body size, changes in which could explain some of the variability in animal weights.

### Gene ontology (GO) term analysis reveals a link of candidate genes to the central nervous system (CNS)

Next, we used GO term analysis to further explore the functions of the genes identified in the GWASs as linked to animal weight. GO term analysis of the 23 genes linked by the GWAS to weight in the control animals using PANTHER (Mi, Muruganujan, Ebert, et al., 2019; Mi, Muruganujan, Huang, et al., 2019) identified 64 GO terms, many linked to morphogenesis and the CNS (Figure 4A and Table S4). Specifically, the high level GO terms “neuron fate determination”, “regulation of axon extension involved in axon guidance”, “dorsal closure, amnioserosa morphology change”, “Malpighian tubule development”, “anterior/posterior pattern specification”, “open tracheal system development”, “post-embryonic appendage morphogenesis”, “ imaginal disc-derived appendage morphogenesis”, “imaginal disc morphogenesis”, “cellular component morphogenesis”, and “cell development” suggest that many of the genes identified in the GWAS function in the CNS and/or morphogenesis. GO term analysis of the genes linked to weight in exercise-treated animals also identified several biological processes that are overrepresented (Figure 4A and Table S4). Specifically, the high level GO terms “axon extension”, “axon guidance”, “neuron recognition”, “cell junction assembly”, “anatomical structure homeostasis”, “cell migration”, and “cell communication” were identified as significantly enriched in the gene set. The prevalence of terms related to the CNS in both analyses suggests that the CNS might extend significant control over feeding behavior and animal activity, thus impacting animal weight.

**Figure 4.**
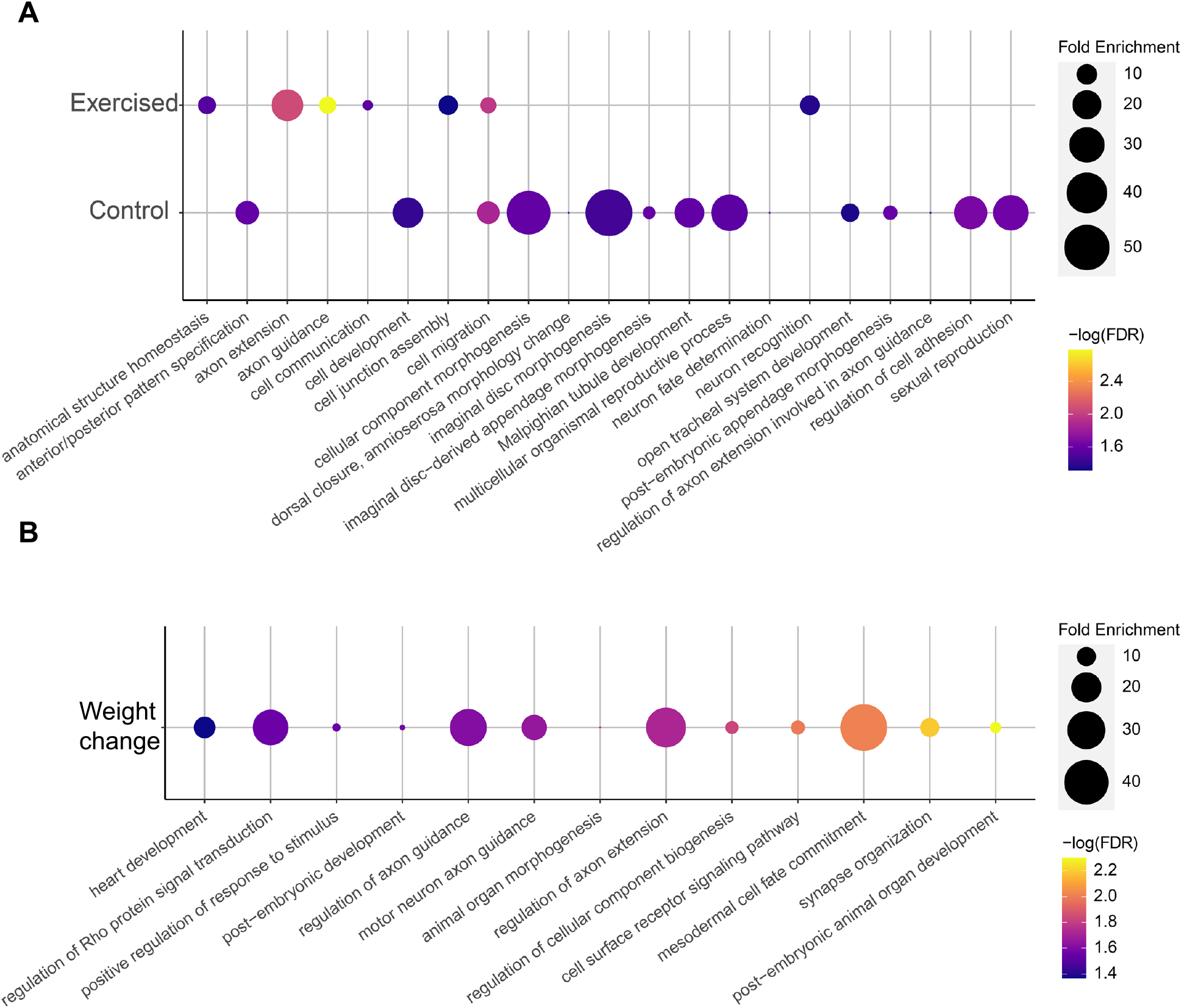
GO term analysis suggests a link between body weight and the CNS. High level GO terms significantly overrepresented in the gene set identified by the body weight GWAS analyses are shown, with fold enrichment shown by the size of the circles and the p-value (FDR) for the over-representation test (Fisher’s Exact Test) shown by the color of the circles. Top: Data from exercised animal weight analysis; bottom: Data from exercised animal weight analysis.

### Differences in animal weight are linked to differences in body composition

While the GWAS results suggest a link between animal weight and the CNS, they do not resolve the question if heavier animals are larger than lighter animals – as is typically seen for the two sexes – or if the heavier animals are “denser”, due to differences in body composition. To gain further insights into the origin of the weight differences observed, we investigated the body composition of a selection of five DGRP strains. Using QMR, we determined total weight as well as lean and fat weight (Figure 5). Again, we found that there was significant variation among the five DGRP strains in terms of total animal weight, with females weighing more than males in all strains (ANOVA, sex effect p = 4.93e-14, genotype effect p = 0.000156). There was also variation in terms of percent fat mass, which ranged from approximately 15% to 22%. The percent of fat mass was influenced strongly by sex (Wilcoxon rank test, p = 0.003211), with females having higher body fat levels than males in four of the five strains investigated. The percent of lean mass varied similarly, ranging from approximately 67% to 78%. Again, the percent of lean mass was influenced strongly by sex (Wilcoxon rank test, p = 0.00234), but here, males had higher levels of lean mass than females in four of the five strains investigated. When considering the relationship between total weight and body composition, there was a negative correlation between percent lean mass and body weight, suggesting that animals with lower weights show more lean mass (Pearson’s product-moment correlation r = −0.632081, p = 1.215e-05). On the other hand, there was a positive correlation between percent body fat and total weight, suggesting that animals with higher weights have more body fat (Pearson’s product-moment correlation r = 0.5720437, p = 0.0001151). These data suggest that in addition to weight, body composition differs significantly between the DGRP lines and that body composition might impact total animal weight, with higher levels of body fat being associated with increased weight.

**Figure 5.**
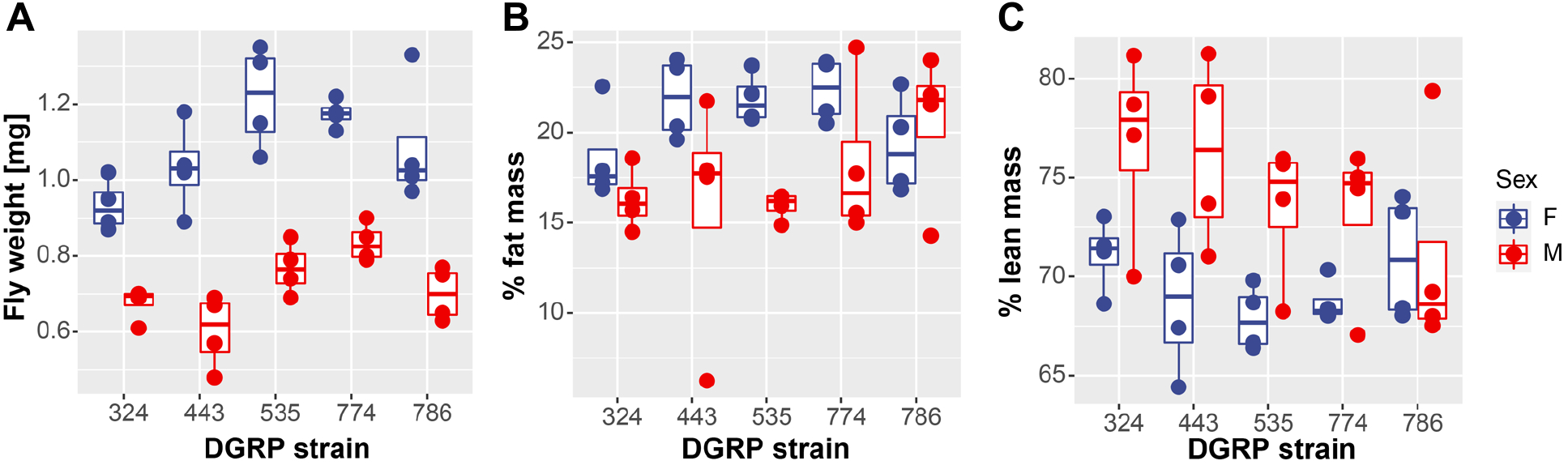
Body composition analysis indicates that heavier animals show higher body fat and less lean mass. Fly weight (**A**), percent fat mass (**B**), and percent lean mass (**C**) as measured by QMR are shown on the Y-axis for five different DGRP strains (X-axis). Data from males is shown in red, data from females in blue. The box plots show the median (horizontal bar) and 25^th^/75^th^ percentiles.

### Heavier flies are typically larger than lighter flies

While the results from the body composition analysis suggest that at least some weight variation among the DGRP strains is due to variation in body composition, they do not rule out that animal size also plays a role. Thus, using the same five DGRP strains included in the body composition analysis, we measured wing size. In Drosophila, wing size correlates well with body size as long as temperature is constant and can be reliably measured (Mirth & Shingleton, 2012; Shingleton et al., 2009). We found that in the five strains investigated, total wing area (measured in pixels) ranged from 143 to 200 in males and 183 to 232 in females. Consistent with their larger weight, females had larger wings in all strains (Figure 6), confirming as expected that females were larger in size than males. In addition to sex significantly impacting wing size (p = 3.76e-07, Kruskal-Wallis rank sum test), genotype also significantly impacted wing size (p = 4.1e-05, Kruskal-Wallis rank sum test). When investigating the relationship between animal weight and wing size, we found a strong positive correlation of 0.845 (Pearson’s product-moment correlation, p = 0.002075; Figure 6). Thus, the analysis of wing size confirms that the animals of heavier DGRP strains are typically larger in size, irrespective of body composition.

**Figure 6.**
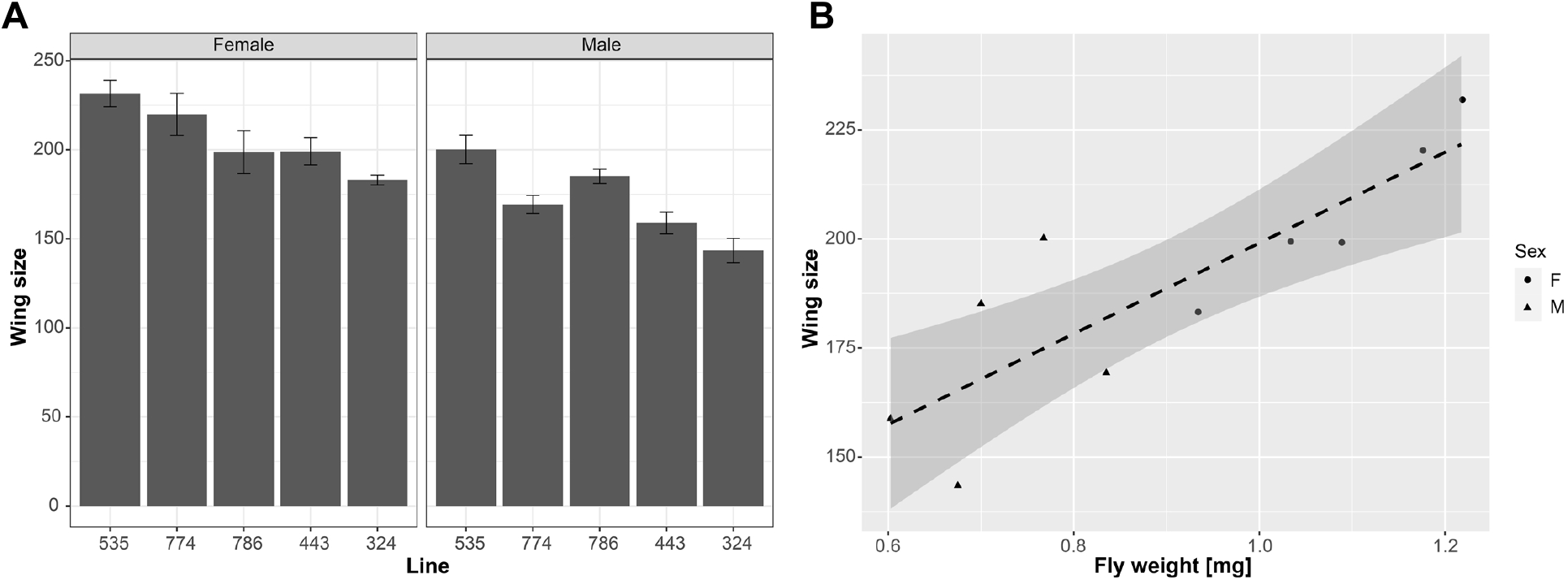
Wing size is strongly correlated with animal weight. **A.** Wing size (Y-axis, measured in pixels) varies significantly in the DGRP population (Lines, X-axis), with females (left) having consistently larger wings than males (right). Error bars: standard errors, n = 10. **B.** Fly weight (X-axis, in mg) is strongly correlated with wing area (Y-axis, measured in pixels). Trend line shown in black (dashed) with standard error. Data from females are circles, data from males are triangles.

### Drosophila weight can change in response to a five-day exercise treatment, and in female flies, this weight change is linked to the level of exercise intensity

A major goal of this study was to determine how animals respond to an exercise treatment. Thus, we compared the weights of animals after the completion of a five-day exercise treatment to the weight of control animals that had been handled in an identical fashion but without an exercise induction (Figure 7). We found that in females, the weight differences ranged from a weight loss of 0.080 mg (line 859) to a weight gain of 0.108 mg (line 727). In males, the weight differences ranged from a weight loss of 0.068 mg (line 374) to a weight gain of 0.079 mg in line 486. Looking at this weight difference in percent of the control animal weight, we saw changes ranging from −6.6% to +11.0% in females and from −8.2% to +11.3% in males. This wide range of weight changes observed in response to the exercise treatment suggests that, just like humans, fruit flies vary in their physiological and behavioral responses to exercise stimulation.

**Figure 7.**
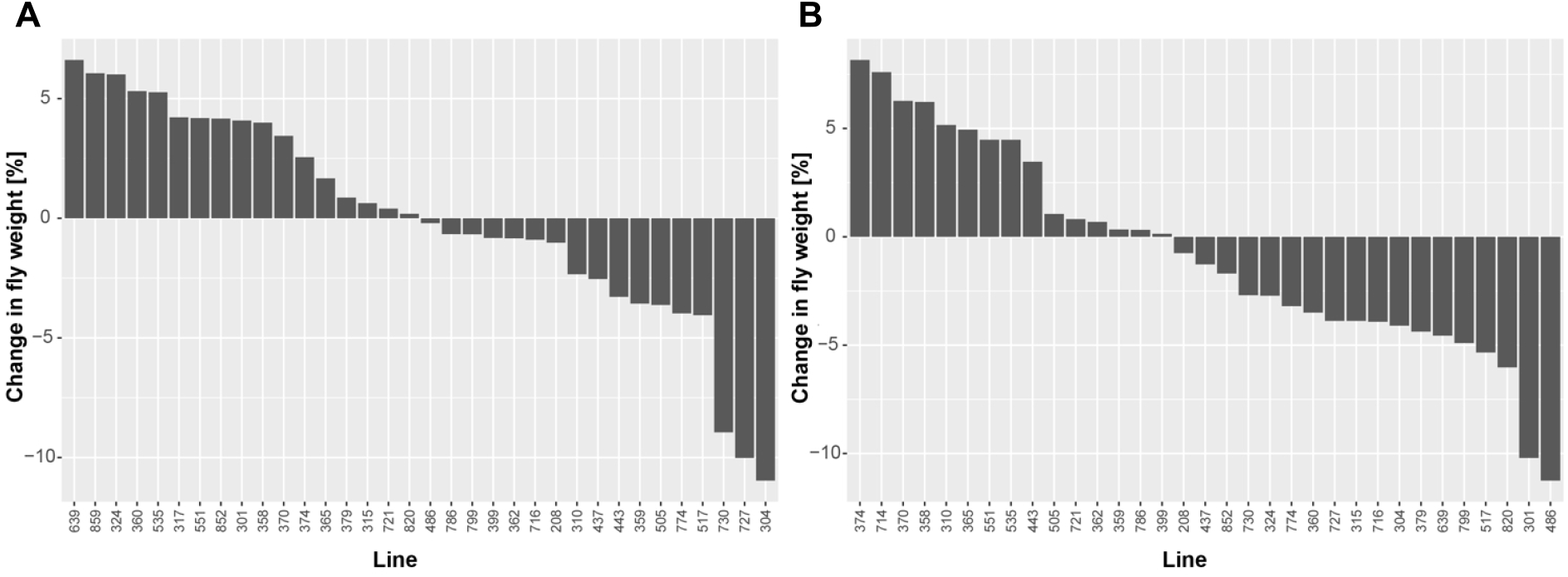
Weight gains as well as losses are seen after exercise treatment. X-axis – DGRP line number; Y-axis – weight change relative to control in % calculated as (control-treatment)/control*100. **A.** Data from females. **B.** Data from males.

One likely reason why these weight changes in response to the exercise treatment are highly variable, might be that the DGRP strains differ in their activity levels during a rotational exercise treatment (Watanabe et al., 2020). To address this possibility, we investigated the link between the change in body weight observed and the intensity of the exercise performed by the animals, using the difference in activity between basal and exercise-induced activity levels as a measure of exercise intensity. In a simplistic model, one would expect that animals that show the greatest change in activity upon exercise treatment would also show the greatest change in body weight. Thus, we calculated the correlation between the change in body weight observed in this study and the previously reported change in activity upon rotational exercise induction (Watanabe et al., 2020). We found that in males, there was no significant correlation between the exercise-related change in animal activity and body weight change after the 5-day exercise treatment (r = −0.193; p-value = 0.3458; Figure S3). Interestingly, in females, we detected a significant correlation between exercise-induced activity change and exercise-induced weight change. The Pearson product moment correlation was 0.503 (p-value = 0.007447), indicating that female animals with the greatest increase in activity level also show the greatest increase in weight in response to the 5-day exercise regime used here. This finding suggests that in female animals, where the rotational exercise treatment leads to a more robust increase in activity levels than in male flies, a physiological response to the exercise in terms of animal weight can be detected.

### A GWAS of the weight change experienced by animals following a five-day exercise treatment reveals a link to signaling

To investigate how genetic factors impact weight change associated with exercise, we carried out a GWAS. We used the data from Figure 7 as basis for the GWAS, again using the DGRP2 webtool (Huang et al., 2014; Mackay et al., 2012). The analysis identified 89 genetic variants linked to exercise-induced weight change, either in males, females, combined sexes, or associated with the difference between sexes (Table S3). These variants represented a total of 52 genes. These genes again included some that are described on FlyBase as being “seize defective” FB2020_03) (FlyBase et al., 2004; Gramates et al., 2017; Thurmond et al., 2019), such as *BHD*, *klar*, *Lar*, *Nlg1*, *Ptp61F*, *pum*, *spz5*, and *stan*. Another gene of note was *mAcon2*, which encodes a mitochondrial aconitase. *mAcon2* is involved in tricarbonic metabolism and has been linked to exercise by work of the Wessells’ lab, reporting increased aconitase levels in exercised flies (Piazza et al., 2009). *α-Man-Ia* is also linked to carbon metabolism; it is an α-mannosidase that glycosylates protein targets and has been linked to longevity (Kerscher et al., 1995; Liu et al., 2009). Overall, there was little overlap between the genetic variants linked to exercise-induced weight change and those variants linked to animal weight, either in the control or exercise-treated animals. There were two genes in common between the results of the weight change GWAS and GWAS for weight in control animals (*Lar* and *Ptp61F*) and three genes shared between the results from the weight change GWAS and the GWAS for weight in exercise-treated animals (*Pka-R2*, *Ptp61F*, and *pum*). Thus, the analyses carried out here suggest that, not surprisingly, different genetic pathways control weight and how an individual’s weight changes with exercise.

To find out more about the genes identified as linked to exercise-induced weight change, we carried out GO term analysis with the PANTHER tool set (Mi, Muruganujan, Ebert, et al., 2019; Mi, Muruganujan, Huang, et al., 2019). This analysis identified a large number of GO terms significantly overrepresented in the candidate gene set (Table S4, FDR < 0.05, Fisher’s Exact test), including 14 high level GO terms (Figure 4B). We again saw terms related to the CNS, such as “regulation of axon extension,” “regulation of axon guidance,” and “motor neuron axon guidance.” A new class of GO terms that did not appear to be involved in the control of weight in general were linked to signaling. These terms included “regulation of Rho protein signal transduction”, “cell surface receptor signaling pathway”, and “positive regulation of response to stimulus”. In addition, a Reactome pathway analysis conducted using PANTHER identified eight pathways significantly overrepresented among the candidate gene set, all of which are signaling pathways (Table S4, FDR < 0.05, Fisher’s Exact test). Thus, the GO term analysis suggests that a variety of signaling pathways are important for weight change in response to exercise, which might indicate that for responses to exercise to occur a variety of molecular pathways mediating signaling between organ systems are needed.

## DISCUSSION

The primary objective of this research was to gain insight into the genetic architecture and molecular pathways controlling animal weight and exercise-induced weight change using the Drosophila model. Our investigation confirmed the previous finding by Jumbo-Lucioni and colleagues that the DGRP population contained significant variation in animal weights (Jumbo-Lucioni et al., 2010). However, despite the fact that a similar number of DGRP lines were sampled in both studies, many overlapping, the animal weights observed in our study represent a wider range. In addition, the heritability for body weight reported here (0.773 and 0.758) is markedly higher than the heritability reported by Jumbo-Lucioni (0.25), possibly due to differences in rearing conditions and food composition (Jumbo-Lucioni et al., 2010). The GWAS results obtained in these two studies also differ, with our study reporting approximately 100 genes linked to animal weight, while Jumbo-Lucioni and colleagues identify over 300 genes, with no overlap between the gene sets (Jumbo-Lucioni et al., 2010). This difference between the two studies also carries to the functional classification of genes associated with the candidate genes discovered. In the study presented here, candidate genes are enriched for GO terms associated with development and morphogenesis, especially as it relates to the CNS. The Jumbo-Lucioni study also notes a link to the CNS, but they find a large number of genes linked to “defense,” which is absent in our analyses (Jumbo-Lucioni et al., 2010). Part of this difference likely is the method used for the GWAS, as our study used the DGRP webtool, which includes the latest annotation of SNPs from the DGRP and corrects for the presence of Wolbachia and some chromosomal rearrangements. This tool was not yet available at the time Jumbo-Lucioni and colleagues carried out their study, and they used a subset of the SNPs available today and included quantitative trait transcripts in addition to SNPs (Jumbo-Lucioni et al., 2010). Thus, it is possible that the “defense” GO term detected by Jumbo-Lucioni and colleagues is due to the lack of correction for Wolbachia infections that are present in some of the DGRP strains. While the Wolbachia infections do not show an impact on weight in our study (p>0.05, ANOVA), such an infection would be expected to impact gene expression and possibly the quantitative trait transcripts. Thus, the differences between the two studies highlight the fact that animal weight is a complex trait that is impacted by many different pathways, the importance of which is dependent on environmental factors and experimental design.

The complexity of the various factors that influence body weight is further illustrated by considering another study using the DGRP, a study investigating body size and the changes in body size induced by changes in temperature (Lafuente et al., 2018). While there clearly is a relationship between body weight and body size, this relationship is not straight forward. In our study of wing size, we find that larger wings are typically associated with heavier flies. However, when we compare our weight data to that of Lafuente and colleagues, who investigated the temperature-dependence of body size by measuring abdomen and thorax size in the DGRP population (Lafuente et al., 2018), we find no relationship between thorax or abdomen size and fly weight (p>0.05 for Pearson’s product moment correlation between thorax size or abdomen size and female fly weight). There is also no overlap between the genes that control abdominal or thorax size or the temperature-associated variation therein and the genes that we detect as candidate genes impacting animal weight. This finding links to studies from the field of allometry in a variety of animals that demonstrate that body parts do not necessarily scale equally with overall body size, thus leading to different proportions of body parts to each other (Mirth et al., 2016; Vea & Shingleton, 2020). If the various body parts then have different densities, which is the case for muscle versus body fat, the body weight does not simply reflect size. Instead, as the QMR experiment presented here suggests, the body weight variation in the DGRP population is due to a combination of the effects of size and body composition, i.e. density of tissues.

With regard to the exercise-induced weight change, the large amount of variation seen in the response, ranging from weight gain to weight loss, is consistent with what has been observed in mice as well as humans (Hiramatsu & Garland, 2018; Kelly et al., 2014; King et al., 2017; Pickering & Kiely, 2019; Solomon, 2018; Stephens & Sparks, 2015; Vellers et al., 2018). The fact that female flies show a clear correlation between the amount of exercise performed and the exercise-induced weight change is a promising result, that will require further study. As the weight change is an increase in weight, and because females typically have a higher amount of body fat, it is tempting to speculate that exercise induced a shift from fat mass to muscle mass in the females that showed the biggest increase in activity levels with exercise. The DGRP population is an ideal resource to further test this model, as it includes lines like line 786, which does not show a difference in body fat level between males and females. As the fly’s size remains unchanged due to the constraints of its exoskeleton, all change in exercise-induced weight change are due to a shift in the density of the animal, most likely shifts between body fat and lean mass. Thus, flies provide an exciting opportunity to dissect the molecular pathways that control this shift and to explore additional exercise treatments to identify optimal parameters.

A final important take-away from this study is the finding that the CNS plays a significant role in controlling animal weight, and that signaling is particularly important for the exercise-induced weight change. This finding is consistent with the results from Jumbo-Lucioni and colleagues that also find a link between animal weight and the CNS in their study (Jumbo-Lucioni et al., 2010). At this point, a variety of studies from diverse systems hint at the importance of the CNS in controlling animal weight (Goodarzi, 2018; Kim et al., 2018; Ryan et al., 2012). This influence of the CNS is mediated by a variety of pathways and includes hormonal influences, impacts via nutrient sensing, and behavioral modifications initiated by the CNS. Thus, many of these pathways include long-range signaling cascades that require the communication between various tissues and body parts (Kim et al., 2018). This type of long-range communication between the CNS and the other parts of the body is an intriguing avenue of research, and its role in the regulation of exercise responses is a largely unexplored area.

In summary, this study reveals the complex genetic architecture that influences animal weight as well as exercise-induced weight change. We demonstrate that genetic background is an important determinant of how individuals respond to exercise and that it needs to be taken into account to design optimal exercise regimes and exercise-based medical interventions. Personalized exercise treatments will be needed to address the obesity epidemic and to allow every individual to reap the maximum benefits from exercise. The Drosophila system with its excellent genetic tools and the ability to study large sample sizes provides unique opportunities to further investigate the roles of long-range CNS communication and body composition shifts in the response to exercise to identify trackable factors to be prioritized for further study in mammals.

## MATERIALS AND METHODS

### Fly Husbandry

All fly strains used in this study are part of the Drosophila Genetics Reference Panel (DGRP, Table S1) (Huang et al., 2014; Mackay et al., 2012). They were obtained from the Bloomington Drosophila Stock Center or Dr. L. Reed (University of Alabama). The animals were reared in a temperature-, humidity-, and light-controlled incubator (25°C; 70% humidity; 12 h : 12 h light : dark cycle; lights on: 7:00 h – 19:00 h) in vials on media consisting primarily of cornmeal, agar, molasses, and yeast, with propionic acid and tegosept added as antifungal agents (Mendez et al., 2016). Population densities were controlled by starting vials with ten virgin females and seven males. Animals used in the body composition and wing size analyses were grown on Jazzmix media (Thermo Fisher Scientific, Waltham, MA).

### Exercise Regime

The exercise treatment used in this study was a five day exercise regime, and the study also included control animals that did not receive an exercise treatment. Flies were exercised on the TreadWheel as described by Mendez and colleagues, following the protocol used in “Study B” from The University of Alabama at Birmingham (Mendez et al., 2016). Briefly, 3-5 day old virgin flies were treated in groups of 20 animals, five vials for each sex-genotype-treatment combination, in vials containing one inch of media. The treated animals exercised for two hours per day for five days, while vials of control animals were placed on a platform attached to the TreadWheel in order for the animals to experience the same vibrations and noise generated by the TreadWheel without the rotational stimulation. The flies were loaded onto the TreadWheel at 11:00 h each day for a one-hour acclimatization period. At 12:00 h, rotation was initiated at 4 rotations-per-minute, and the flies were exercised for two hours. Following each exercise period, the vials were unloaded from the TreadWheel and returned to the climate-controlled incubator until the next exercise period. Upon completion of the five day exercise regime, vials of flies were stored in the incubator until follow-up assays were performed the next day.

### Mass Assay

On the day following the completion of the exercise treatment, all animals were subjected to a climbing assay protocol adapted from the rapid iterative negative geotaxis assay (RING) to assess exercise response (Gargano et al., 2005). Immediately following these climbing assays, flies were anesthetized with CO_2_ and separated into groups of 10 by genotype/sex/treatment combination, placed into microcentrifuge tubes, and weighed on a microscale. Typically, sample size was ten (ten microcentrifuge tubes containing ten flies each) per genotype/sex/treatment combination.

### Genome-wide Association Studies

GWASs were carried out for the trait “average control weight” and “average treatment weight” using the means per fly weights for each genotype*sex combination. Genetic variants linked to the two phenotypes were identified using the GWAS webtool developed for the DGRP by Mackay and colleagues (http://dgrp2.gnets.ncsu.edu/) (Huang et al., 2014; Mackay et al., 2012). Based on the approach taken in other studies of the DGRP and qq-plot analysis, a nominal p-value cutoff of 10^−5^ was used to identify candidate genes for further consideration (Arya et al., 2015; Campbell et al., 2019; Dembeck et al., 2015; Jordan et al., 2012; Mackay et al., 2012; Swarup et al., 2013; Weber et al., 2012).

### Gene ontology (GO) term analysis

GO term analysis was carried out using the PANTHER (Protein ANalysis THrough Evolutionary Relationships; version 15.0) Classification System (Mi, Muruganujan, Ebert, et al., 2019; Mi, Muruganujan, Huang, et al., 2019). A false discovery rate (FDR; p<0.05) was used for multiple testing correction in the overrepresentation test (Fisher’s Exact test).

### Body composition analysis by quantitative magnetic resonance (QMR)

Body composition was analyzed by QMR at the UAB Small Animal Phenotyping Core as described previously (Mills et al., 2018). Briefly, virgin flies were collected and processed at 0-4 days of age. Ten flies were used per replicate, and four replicates per sex and genotype. Groups of flies were moved into the biopsy tube and scanned using the biopsy setting with nine primary accumulations with an EchoMRI 3-in-1 machine (Echo Medical Systems, Houston, TX).

### Wing size analysis

Wings were removed from animals grown at the same time and under the same conditions as the animals used for the QMR analysis described above. The wings were removed from the animals using dissecting scissors, temporarily stored in 1x TBST, and then placed on a microscope slide. After adding a cover slip, the wings were imaged with an AZ100 microscope (Nikon) with 4x magnification. The resulting images were processed using FijiWings, obtaining the total wing area as the number of pixels using the “lasso” tool (Dobens & Dobens, 2013).

### Statistical Analyses

All statistical analyses were performed using R (Team, 2018). The restricted maximum likelihood (REML) approach for mixed models and the VCA package (Schuetzenmeister & Dufey, 2020) were utilized to estimate variance components with the following model: Y = μ + S + L + SxL + ε (L, line, random; S, sex, fixed). The phenotypic variance (σ_P_^2^) includes genetic (σ_G_^2^) and environmental (σ_E_^2^) with the genetic variance including both the variance due to line and the sex by line interaction (σ_G_^2^ = σ_L_^2^ + σ_L*S_^2^) and the environmental variance defined as the within line variance (σ_P_^2^; σ_P_^2^ = σ_G_^2^ + σ_E_^2^). Broad sense heritability (H^2^) was calculated H^2^ = σ_G_^2^/σ_P_^2^. Coefficients of genetic and environmental variance are CV_G_= 100σ_G_/mean and CV_E_= 100σ_E_/mean. The cross sex genetic correlation was r_MF_ = σ_L_^2^/(σ_L_^2^ + σ_L*S_^2^), and genetic correlation was r_g_ = σ_L_^2^/sqrt(σ_L_^2^(female data)* σ_L_^2^(male data)).

Analyses of variance (ANOVA) included “sex” and “genotype” as factors, as well as their interaction. If the interaction was not statistically significant, it was removed from the final model.

## ACKNOWLEDGEMENTS

We would like to thank the many Riddle lab undergraduates that contributed to the success of the project, particularly, C. Gordon and M. Azar. M. Momeni and J. Favors assisted in the QMR and wing size data collection. We thank Drs. B. Lamos (Harvard University), K. Maggert (University of Arizona), and C. Queitsch (University of Washington) for stimulating discussions regarding the genetic contributions to animal weight. In addition, we thank the members of the Riddle lab for support, comments on the manuscript, and thoughtful discussions. We thank Dr. L. Reed (University of Alabama) for providing many of the DGRP stocks. Stocks obtained from the Bloomington Drosophila Stock Center (NIH P40OD018537) were used in this study. The UAB Small Animal Phenotyping Core is supported by the NIH Nutrition & Obesity Research Center P30DK056336, Diabetes Research Center P30DK079626 and the UAB Nathan Shock Center P30AG050886A.

## AUTHOR CONTRIBUTIONS

LPW participated in the design of the study, carried out the data collection and preliminary data analyses, and critically revised the manuscript; NCR conceived of the study, designed the study, coordinated the study, carried out the final data analyses, and drafted the manuscript. All authors gave final approval for publication and agree to be held accountable for the work performed therein.

## SUPPLEMENTAL FIGURES AND TABLES

**Figure S1.**
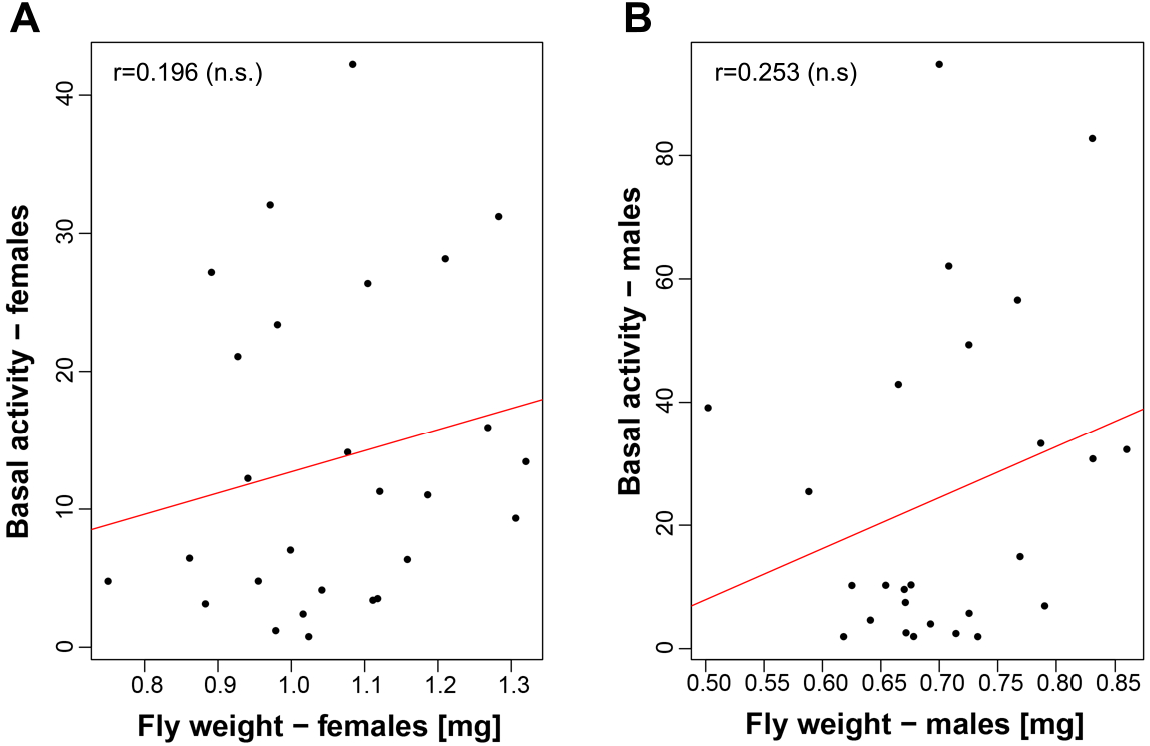
There is no clear relationship between body weight and basal activity levels in the study population. Body weight per fly in mg (X-axis) is plotted against basal activity levels (Y-axis; from (Watanabe et al., 2020)), with data from females shown in **A** and data from males in **B**. For both correlations, the p-value is not significant (p>0.05).

**Figure S2.**
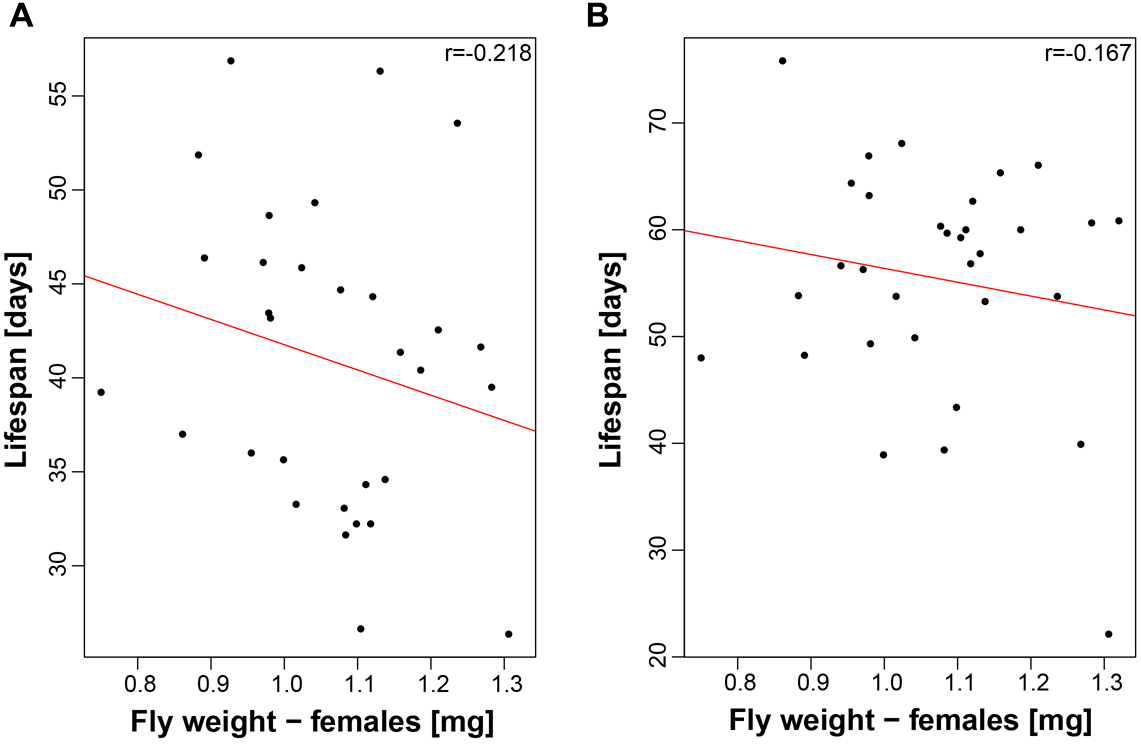
There is no clear relationship between body weight and lifespan in females of the study population. Body weight per fly in mg (X-axis) is plotted against lifespan (Y-axis). Lifespan data in **A** are from Durham and colleagues (Durham et al., 2014), while lifespan data in **B** are from Ivanov and colleagues (Ivanov et al., 2015). For both correlations, the p-value is not significant (p>0.05).

**Figure S3.**
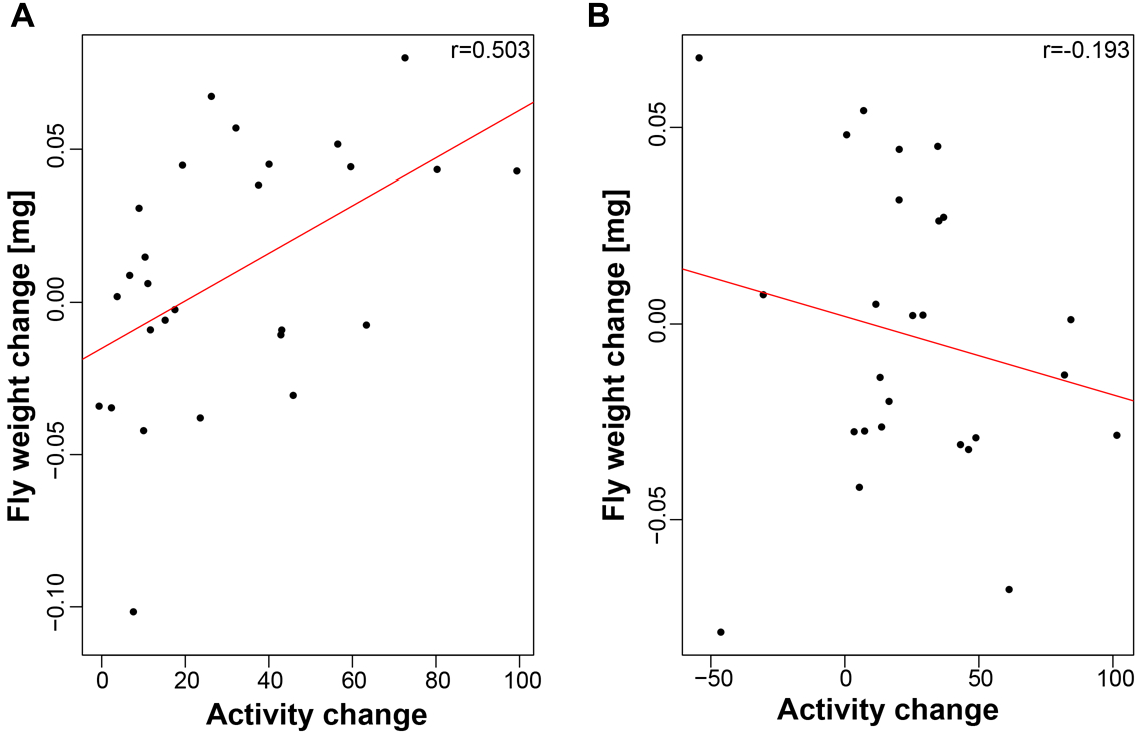
In females, the increase in overall activity associated with an exercise treatment is correlated to the weight change. The change in activity induced by a rotational exercise treatment over baseline (X-axis) is plotted against the change in body weight in mg (Y-axis). Data from females are shown in **A**, displaying a significant positive correlation (p = 0.007447), while data from males in **B** reveal no significant correlation (p = 0.3458).

**Supplemental Table S1. Raw data used in this study.** This file includes a list of all DGRP strains that are part of this study.

**Supplemental Table S2. Quantitative genetics parameters.** Various quantitative genetics parameters such as heritability are reported for weight in the control and treated animals.

**Supplemental Table S3. GWAS results.** This table includes the GWAS output for and information about genetic variants detected as significant in the GWASs.

**Supplemental Table S4. GO term results.** GO term results from the PANTHER database are reported for the genes identified in the GWAS for weight in control animals, in the GWAS for weight in the exercised animals, and in the GWAS for weight change with exercise treatment.

**Supplemental Table S5. QMR data.** This table includes the raw data from the QMR study presented in Figure 6.

**Supplemental Table S6. Wing size data.** This table includes the raw wing size data presented in Figure 7.

