## Supplemental Table 2 for "GWAS reveal a role for the central nervous system in regulating weight and weight change in response to exercise"

**Table S2. Quantitative genetics parameters.**

| Parameter | Symbol | Weight - control | Weight - treated |
| --- | --- | --- | --- |
| Mean | μ | 0.888 | 0.887 |
| Genetic variance | σ_G_^2^ | 0.011 | 0.012 |
| Genetic standard deviation | σ_G_ | 0.107 | 0.109 |
| Environmental variance | σ_E_^2^ | 0.003 | 0.004 |
| Environmental standard deviation | σ_E_ | 0.058 | 0.062 |
| Phenotypic variance | σ_P_^2^ | 0.015 | 0.016 |
| Phenotypic standard deviation | σ_P_ | 0.121 | 0.125 |
| Heritability | H_2_ | 0.773 | 0.758 |
| Coefficient of genetic variation | CV_G_ | 12.001 | 12.305 |
| Coefficient of environmental variation | CV_E_ | 6.498 | 6.952 |
| Cross-sex genetic correlation | r_MF_ | 0.771 | 0.727 |
| Genetic correlation | r_g_ | 0.883 | 0.863 |
